# Mechanical stimulation promotes enthesis injury repair by mobilizing Prx1^+^ cells via ciliary TGF-β signaling

**DOI:** 10.1101/2021.09.21.461317

**Authors:** Han Xiao, Tao Zhang, Changjun Li, Yong Cao, Linfeng Wang, Huabin Chen, Shengcan Li, Changbiao Guan, Jianzhong Hu, Di Chen, Can Chen, Hongbin Lu

## Abstract

Proper mechanical stimulation can improve rotator cuff enthsis injury repair. However, the underlying mechanism of mechanical stimulation promoting injury repair is still unknown. In this study, we found that Prx1^+^ cell was essential for murine rotator cuff enthesis development identified by single-cell RNA sequence and involved in the injury repair. Proper mechanical stimulation could promote the migration of Prx1^+^ cells to enhance enthesis injury repair. Meantime, TGF-β signaling and primary cilia played an essential role in mediating mechanical stimulation signaling transmission. Proper mechanical stimulation enhanced the release of active TGF-β1 to promote migration of Prx1^+^ cells. Inhibition of TGF-β signaling eliminated the stimulatory effect of mechanical stimulation on Prx1^+^ cell migration and enthesis injury repair. In addition, knockdown of *Pallidin* to inhibit TGF-βR2 translocation to the primary cilia or deletion of *IFT88* in Prx1^+^ cells also restrained the mechanics-induced Prx1^+^ cells migration. These findings suggested that mechanical stimulation could increase the release of active TGF-β1 and enhance the mobilization of Prx1^+^ cells to promote enthesis injury repair via ciliary TGF-β signaling.

## Introduction

Rotator cuff (RC) tear is common in modern sports activity, which often causes persistent shoulder pain and dysfunction.^[1]^ Surgical repair has been a well-established and commonly accepted treatment for severe RC tear, especially when conservative treatment fails.^[2,3]^ It has been reported that approximately 450,000 patients were accepted RC repairs surgery annually in the United States.^[4]^ However, the results of surgical repair are not always satisfactory.^[5]^ Previous studies have shown structural failure of the RC repair ranging from 16% to 94%, with poor outcomes associated with failed microstructure regeneration of RC enthesis.^[4,6,7]^ Histologically, the RC enthesis has been consisted of 4 distinct yet continuous tissue layers: bone, calcified cartilage, uncalcified cartilage, and tendon. This kind of structure can subtly transfer force from muscle to bone, while enthesis functional injury repair remains an insurmountable challenge in sports medicine.^[8]^ Therefore, how to promote regeneration of the enthesis is an urgent problem for clinicians.

Moderate mechanical load is essential for enthesis development and maintenance.^[9]^ Meanwhile, the clinical application of mechanobiological principles following enthesis injury forms the basic rehabilitation protocols.^[10,11]^ However, there is a debate about the initiating time and strength of mechanical stimulation during enthesis healing procedure.^[12]^ In clinical treatment, a traditional rehabilitation program after RC repair has been suggested to delay mechanical exercise (immobilization for about six weeks),^[13]^ while an accelerated protocol suggests that an immediate exercise with limited range of motion would be better for tendon healing.^[14]^ Zhang et al adopted treadmill training at postoperative day seven on murine enthesis injury repair model and found that mechanical stimulation could improve enthesis fibrocartilage and bone regeneration and obtained better mechanical parameters.^[15]^ Although we know that there is a correlation between appropriate mechanical stimulation and high quality of enthesis healing, the uncovering mechanism is poorly understood. Revealing the cellular and molecular processes of enthesis healing with mechanical stimulation after surgical repair will allow clinicians to implement preventative interventions and prescribe proper therapeutics to improve clinical outcomes.

During the wound regeneration procedure, soluble inflammatory mediators bind to cell surface or cytoplasmic receptors, and lead to recruitment of immune cells, stem cells and tissue-resident cells by activating signaling cascades.^[16]^ As we known, stem cells are essential for enthesis regeneration. *Prx1* is a paired-related homeobox gene that is expressed in undifferentiated mesenchymal stem cells in the developing limb buds.^[17]^ A previous study showed that *Prx1* transgene marked osteochondral progenitors in the periosteum and played an essential role in skeletal development.^[18]^ Mice lacking *Prx1* transgenes would show craniofacial defect, limb shortening, and incompletely penetrant Spina bifida.^[19]^ Considering Prx1^+^ cells are indispensable stem cell lineage for the musculoskeletal system, we want to specificlly reveal the role of Prx1^+^ cells in enthesis injury repair and uncover the mechanism of mechanical stimulation on improving enthesis injury repair in this study.

Primary cilia is an antenna-like sensory organelle based on immotile microtube, and present on nearly every cell type, including mesenchymal stem cells, endothelial cells (ECs), epithelial cells, fibroblasts, and other cells in vertebrates.^[20–23]^ Primary cilia contains a distinct subset of receptors and other proteins, which make it a sophisticated signaling center functioning as mechanosensor and chemosensation.^[24–27]^ Previous study also found that translocating receptors to the primary cilia could enhance the signaling transmission.^[28]^ Defect or dysfunction of primary cilia could lead to severe disorders of the body, which is known as ciliopathies, such as polycystic kidney disease, primary ciliary dyskinesia, retinopathies, combined developmental deficiencies, and other sensory disorders.^[29–33]^ Genetic deletion of *IFT88*, an encoded protein closely associated with cilia formation and maintenance, could decrease the load-induced bone formation.^[34]^ At the same time, the primary cilia is a hub for transducing biophysical and hedgehog signals to regulate tendon enthesis formation and adaptation to loading.^[35,36]^ Therefore, we wonder if primary cilia also plays an important role in mechanical stimulation signal transmission during enthesis healing procedure.

In this study, we first revealed the characteristics of Prx1^+^ cells in the developing enthesis by single cell RNA sequencing (scRNA-seq) and examined the dynamic pattern of Prx1^+^ cells in murine RC enthesis at different ages. Then, we used the murine enthesis injury model to find out the mechanism of proper mechanical stimulation on stimulating Prx1^+^ cell migration and enthesis injury repair. Our data demonstrated that appropriate mechanical stimulation could increase the release of active TGF-β1 and enhance mobilization of Prx1^+^ cells to promote enthesis injury repair via ciliary TGF-β signaling.

## Results

### scRNA-seq analysis reveals the cell populations in the developing enthesis

To determine the cellular composition of the developing enthesis, we isolated and sequenced transcriptomes of individual live CD45^−^Ter119^−^ cells from the murine enthesis (from the fully patterned limb at E15.5 to early maturity of enthesis with appearance of a modest 4-zone structure at P28) based on 10× Genomics system (Fig. 1a, b). After sequencing and data quality processing, we got high-quality transcriptomic data from 21532 single-cells, including 8919 E15.5 cells, 7489 P7 cells, and 5124 P28 cells. The single cell RNA sequencing (scRNA-seq) data had high read depth for most of the single cell samples (Figure S1, Supporting Information). We carried out unbiased clustering analysis for all single-cells and identified 23 major cell populations in the developing enthesis by Seurat analysis (Fig. 1c). Through differential gene expression analysis, we annotated 15 clusters into distinct cell types or states based on the expression of genes uniquely or in combinations represented individual cluster identities (Fig. S2). Feature plot showed the canonical marker genes, which enriched in seven enthesis related clusters: BMSCs, Fibroblastic cells, Tenocytes, Chondrocytes, Proliferative stromal cells, Osteoblast/Osteocyte (OB/Ocy), Lepr^+^ cells (Fig. 1d). Cell fraction related to enthesis development showed that the rate of OB/Ocy and Lepr^+^ cells increased significantly in P7 and P28, which was consistent with the enthesis mineralization procedure. At the same time, chondrocytes were high in E15.5, P7 and decreased significantly in P28(Fig. 1e). These data suggested that the maturation of enthesis was higly correlated with osteochondrogenesis procedure.

**Figure 1.**
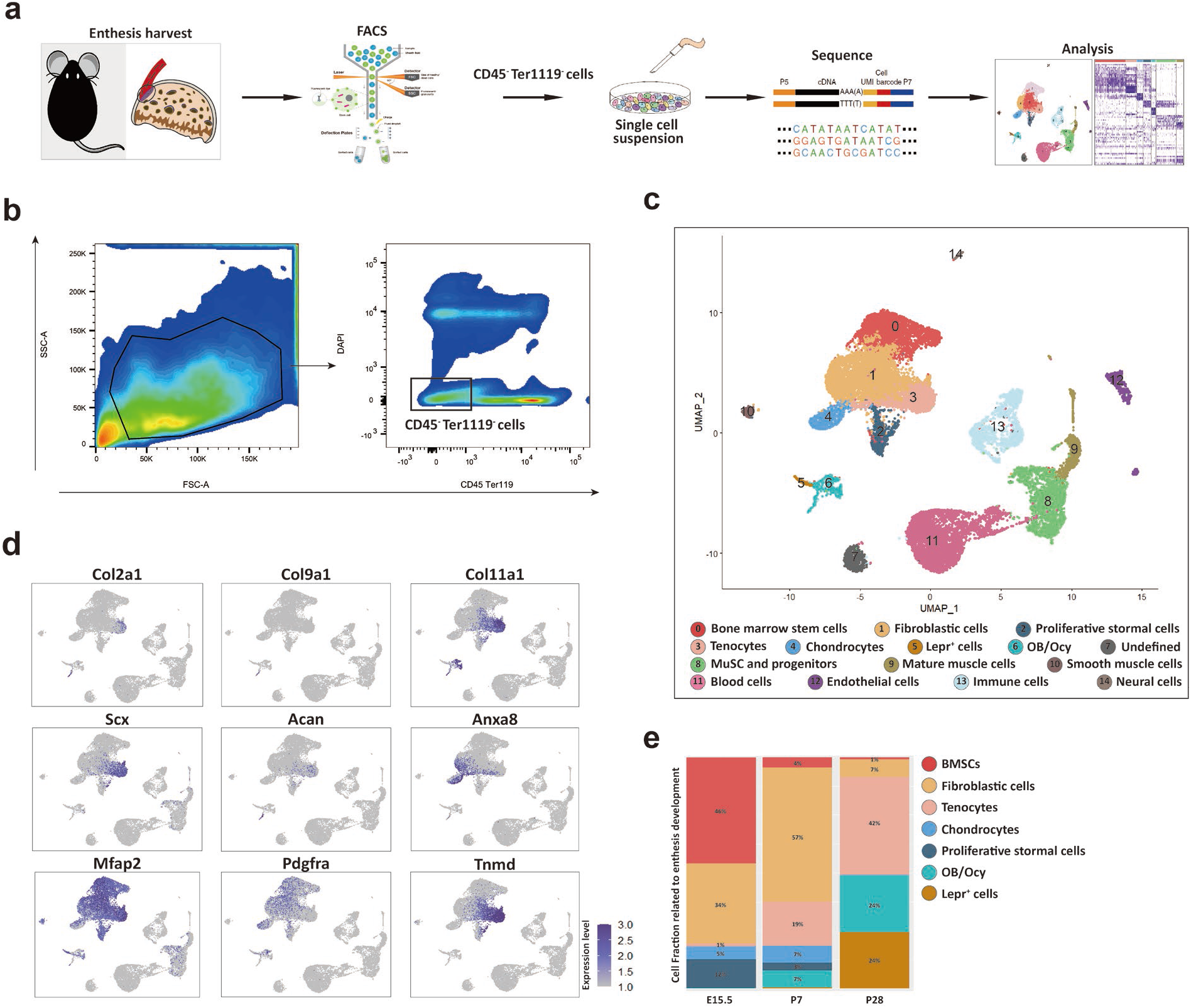
scRNA-seq analysis reveals the cell populations in the developing enthesis. (a) The flow chart of scRNA-seq analysis. (b) Isolation of CD45^−^Ter119^−^ cells by FACS. (c) All cell clusters visualized with uniform manifold projection (UMAP). (d) Feature plot of canonical marker genes enriched in clusters defines enthesis related clusters. (e) enthesis development related cell composition at E15.5, P7, and P28.

### scRNA-Seq distinguishes Prx1^+^ cells during enthesis development

To examing the role of of Prx1^+^ cells in enthesis development, 5607 Prx1^+^ cells from mouse enthesis (E15.5, P7, and P28) were analyzed and were grouped into five distinct clusters: BMSCs, Fibroblastic cells, Tenocytes, Chondrocytes, Lepr^+^ cells. *Prx1* expression was relatively high in E15.5 and P7, while decreased significantly in P28 (Fig. 2a). Pseudotime ordering of Prx1^+^ cells from the enthesis related five clusters were reconstructed by Monocle, an unsupervised algorithm (Figure 2b). The trend of reconstructed trajectory was consistent with the time point (Figure 2c upper panel), which could represent the temporal (stem/progenitor and teno/osetochondro lineage) relationships during the development of enthesis. The reconstructed trajectory tree colored by clusters shows some overlap along the pseudotime (Figure 2c lower panel). These results indicated that Prx1^+^ cells were highly involved in enthesis development via differentiating into tenocytes, osteoblasts/osteocytes or chondrocytes. To further analyze differential gene expression of Prx1^+^ cells in E15.5, P7, and P28, GO enrichment analysis was performed and representative GO terms in represented biological processes were illustrated (Figure2d). The results showed that the ribonucleoprotein complex biogenesis and assembly activities were significantly upregulated in E15.5, while extracellular matrix organization activities in P7 and ossification activities in P28. These data suggested that Prx1^+^ cells could be the potential reliable progenitors for enthesis development.

**Figure 2.**
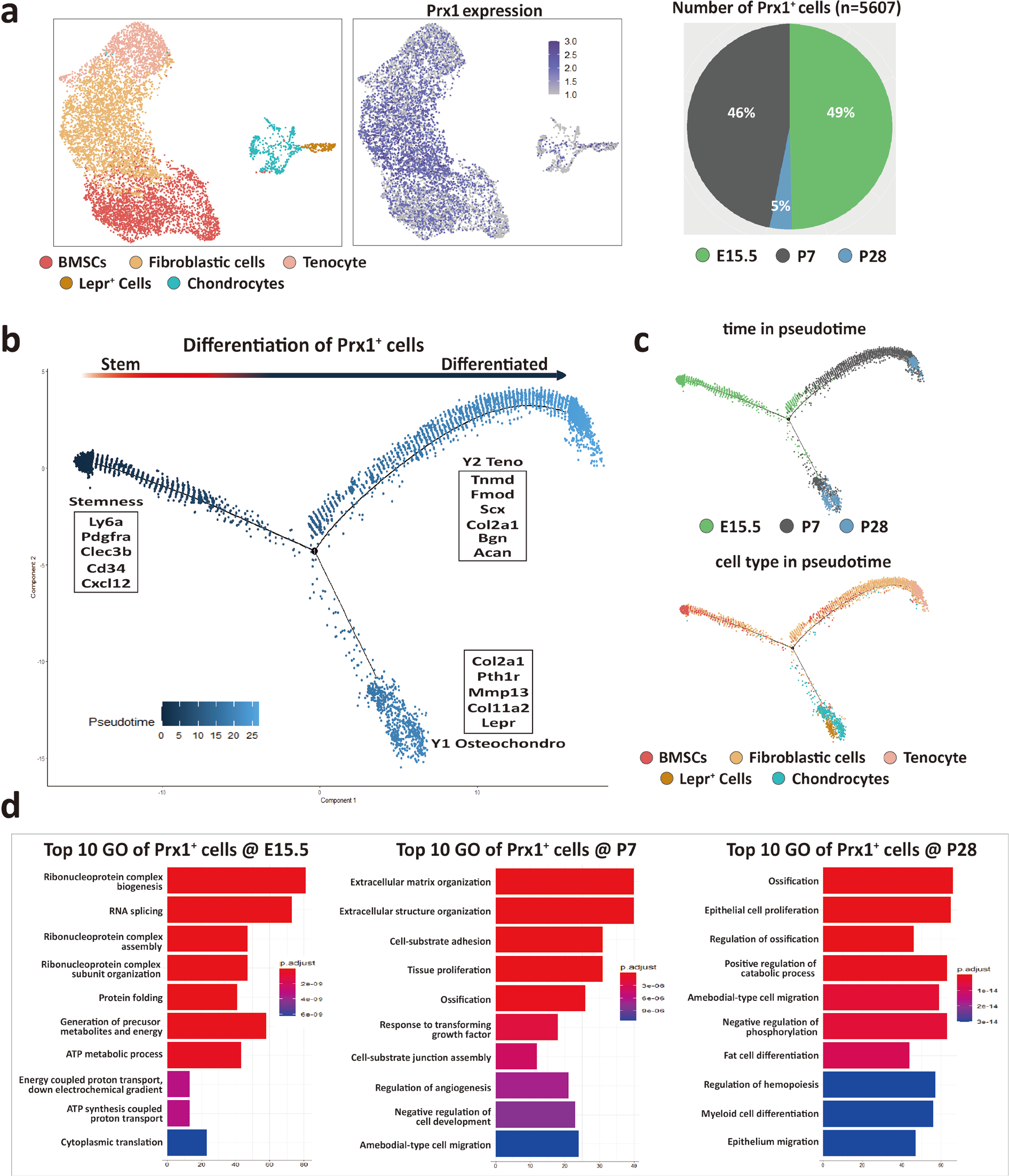
scRNA-Seq distinguishes Prx1^+^ cells during enthesis development. (a) 5607 Prx1^+^ cells from mouse supraspinatus enthesis (E15.5, P7, and P28) were grouped into five distinct clusters (colors indicated). Each point represents an individual cell; Right panel shows the expression level of *Prx1* within enthesis -related clusters. (b) Differentiation trajectory of enthesis -related cells constructed by Monocle and was colored by pseudotime order. Branches on the 2D trajectory tree are indicated as tenogenic branch (Y1) and osteochondrogenic branch (Y2). (c) Uper panel was colored by real time-point and lower panel was colored by cell clusters, respectively. (d) The enriched GO terms (biological processes) of differentially expressed genes in enthesis development related *Prx1^+^* cells at E15.5, P7, and P28, respectively.

### Lineage tracing of Prx1^+^ cells in murine rotator cuff enthesis development and injury repair

To understand the dynamic pattern of Prx1^+^ cells in rotator cuff enthesis, we performed immunostaining of the murine humeral head, using *Prx1CreER-GFP* transgenic mice at P7, P28, and P56. We found that active Prx1^+^ cells (high expression of GFP) were abundant on the peripheral humeral head in young mice and decreased markedly during late adulthood (Figure 3a). At the enthesis, we found active Prx1^+^ cells were present during the early postnatal period, while they decreased significantly with age, then, disappeared and were confined within the perichondrium at adulthood (Figure 3b, c). To investigate the degree of Prx1^+^ cells participating in enthesis development at a different age, we generated *Prx1CreER*; *R26R-tdTomato* mice to permanently label the cells coming from Prx1^+^ cell pool. We respectively injected a single dose of tamoxifen (100 mg/kg, i.p.) into 2 weeks, 4 weeks and 8 weeks old *Prx1CreER*; *R26R-tdTomato* mice and performed immunostaining at 12 weeks (Figure 3d). We found that most of the cells in the enthesis originated from Prx1^+^ cells at the 2W-12W group. At the same time, this involvement decreased significantly at the 4W-12W group and disappeared at the 8W-12W group. Prx1^+^ cells participated in the development of enthesis, including the continuous four gradient layer structure: bone, calcified fibrocartilage, uncalcified fibrocartilage, and tendon (Figure 3e, f). These finding suggested that Prx1^+^ cell was a vital subpopulation of mesenchymal stem cells for enthesis regeneration.

**Figure 3.**
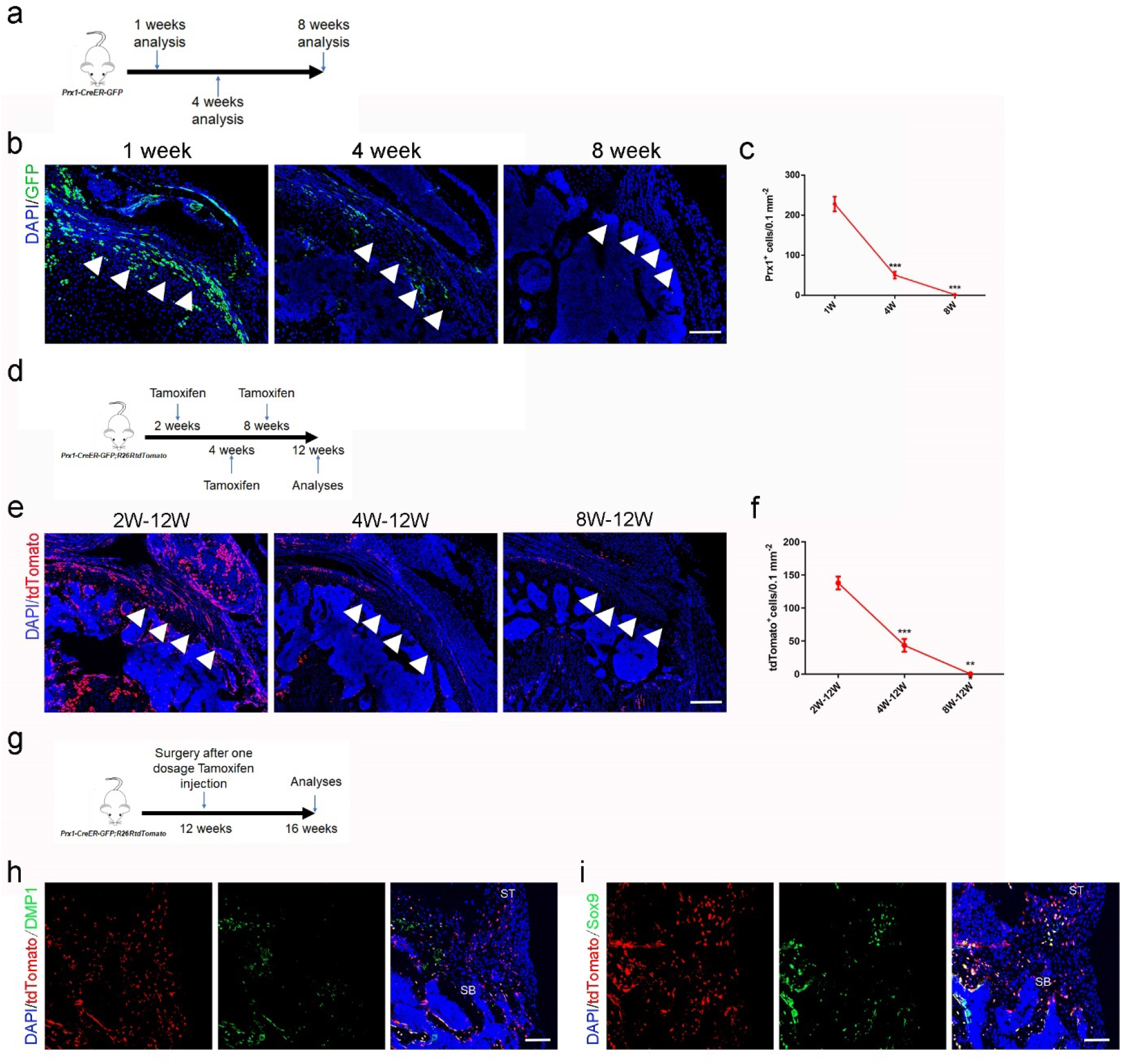
Prx1^+^ cells are involved in rotator cuff enthesis development and injury regeneration. (a) Schematic diagram of *Prx1CreER-GFP* mice, which were sacrificed at 1, 4, and 8 weeks after surgery for immunofluorescent analysis. (b ) Representative immunofluorescent images of GFP staining of active Prx1^+^ cells (green) and DAPI (blue) staining of nuclei in murine humeral head at postnatal 1, 4 and 8 weeks. Scale bars, 200 μm. (c) Quantitative analysis of the number of active Prx1^+^ cells in enthesis. n=6 per group. (d) Schematic diagram of *Prx1CreER-GFP; R26R-tdTomato* mice, which were sacrificed for immunofluorescent analysis at 12 weeks after tamoxifen administration at Postnatal 2, 4, and 8 weeks. (e) Representative immunofluorescent images of tdTomato^+^ cells (Prx1^+^ cells, red) and DAPI (blue) staining of nuclei in murine humeral head at postnatal 12 weeks after injection with tamoxifen respectively at postnatal 2, 4 and 8 weeks. Scale bars, 200 μm. (f) Quantitative analysis of the number of tdTomato^+^ cells in enthesis. n=6 per group. (g) Schematic diagram of *Prx1CreER-GFP; R26R-tdTomato* mice which were received acute enthesis injury and sacrificed for immunofluorescent analysis at 4 weeks after surgery after sigle dose tamoxifen injection. (h) Representative immunofluorescent images of tdTomato^+^ cells (Prx1^+^ cells, red) in murine enthesis at postoperative 4 weeks. Scale bars, 100μm. (i) Quantitative analysis of the number of tdTomato^+^ cells in enthesis. n=6 per group. All data were reported as mean ± SD. The white triangles indicated the area of enthesis. SB, subchondral bone; ST: supraspinatus tendon. *P < 0.05, **P < 0.01, ***P < 0.001.

To verify whether Prx1^+^ cells participated in adult murine enthesis injury healing, *Prx1CreER*; *R26R-tdTomato* mice (12 weeks old) were performed RC injury after injected a single dose of tamoxifen (100 mg/kg, i.p.). At postoperative 4 weeks, mice were sacrificed for immunofluorescence (Figure 1g). We found that Prx1^+^ cells were activated and migrated from the surrounding area to the injury site to participate in the enthesis healing via differentiating into osteocytes or chondrocytes (Figure 3h, i).

### Proper mechanical stimulation improves the enthesis injury repair

To find out if the proper mechanical stimulation could improve enthesis injury repair, the mice began to receive treadmill training at 1 week after enthesis surgery with different treadmill training (0 minutes per day, 10 minutes per day, 20 minutes per day and 30 minutes per day, 5 consecutive days per week). At 4 and 8 weeks after surgery, mice were sacrificed for histology and mechanical test analysis (Figure 4a). We found that treadmill training with 20 minutes per day showed better tissue maturation, collagen arrangement (Figure 4b), higher histological scores (Figure 4c), and more fibrocartilage regeneration (Figure 4d). The best mechanical results of RC have also occurred at the group receiving 20 minutes treadmill training per day (Figure 4e). These results indicated that proper mechanical stimulation could improve enthesis healing, which is correlated with the increased numbers of Prx1^+^ cells.

**Figure 4.**
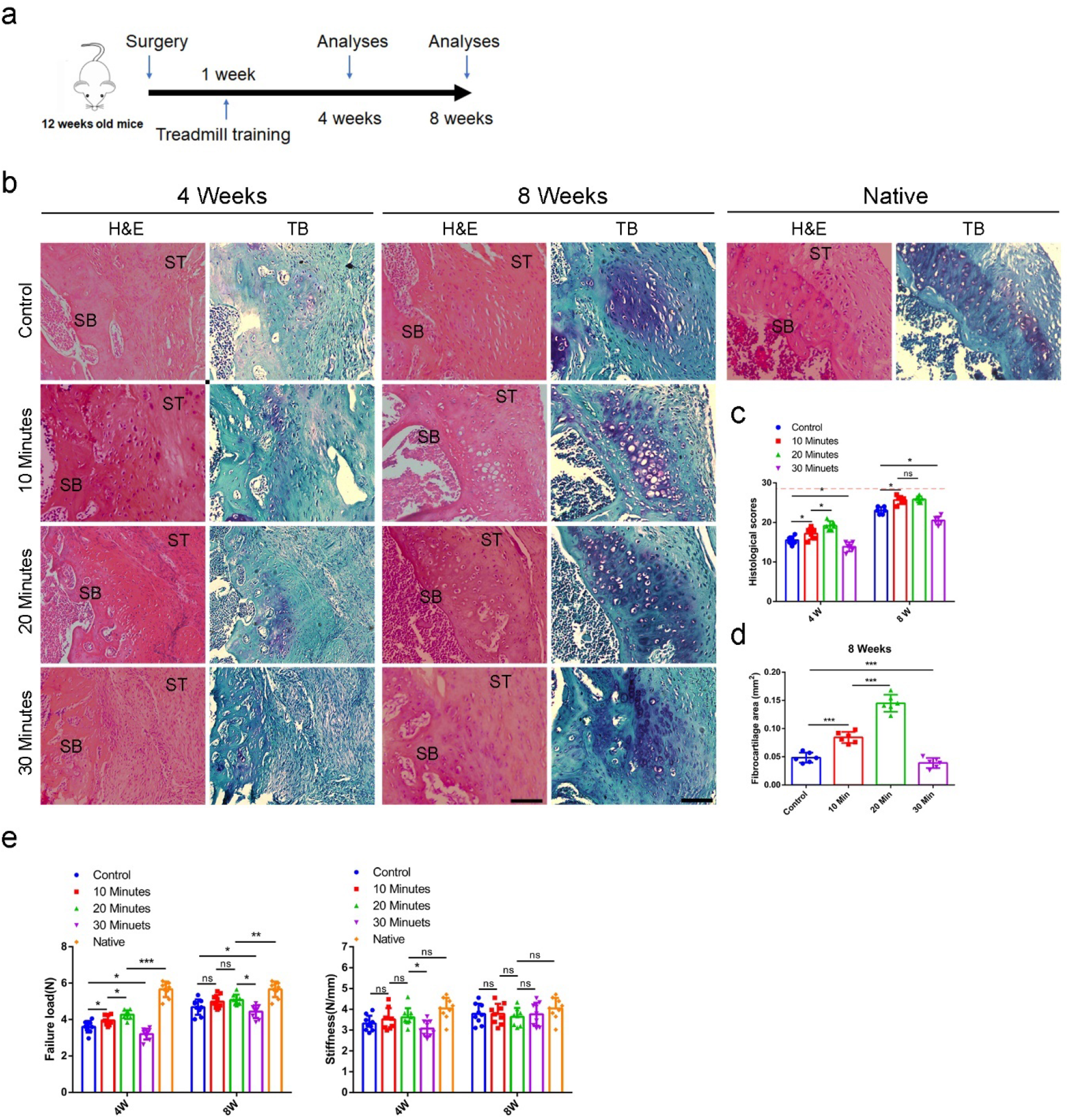
Proper mechanical stimulation could improve the enthesis injury repair. (a) Representative image of H&E and Toluidine blue/Fast green staining of enthesis. Scale bar, 200 μm. (b) Quantitative analysis of H&E score. The red dolt line indicated the perfect histological score of 28. n=6 per group. (c) Quantitative analysis of fibrocartilage thickness. n=6 per group. (d) Quantitative analysis of Failure Load and stiffness. n=9 per group. SB, subchondral bone; ST: supraspinatus tendon. *P < 0.05, **P < 0.01, ***P < 0.001, ns P > 0.05.

### Proper mechanical stimulation mobilize the Prx1^+^ cells to participate in enthesis injury repair

To investigate the potential role of mechanical stimuli on Prx1^+^ cells and the relationship between Prx1^+^ cells number and repair quality, we performed lineage tracing analysis using *Prx1CreER*; *R26R-tdTomato* mice. After receiving enthesis injury repair surgery followed with a single dose of tamoxifen (100 mg/kg, i.p.), the mice started to receive different treadmill training (0 minutes per day, 10 minutes per day, 20 minutes per day, and 30 minutes per day, 5 consecutive days per week) at 1 week after surgery (Figure 5a). We found that Prx1^+^ cells were absent at the enthesis in adult mice (Figure 5b). Prx1^+^ cells could migrate from the nearby tissue to the healing area at 2 weeks after surgery (Figure 5b). The 10-minutes and 20-minutes treadmill training could significantly enhance the migration of Prx1^+^ cells to the healing area compared with the group without treadmill training. Excessive treadmill training decreased the migration of Prx1^+^ cells to the healing area (Figure 5c).

**Figure 5.**
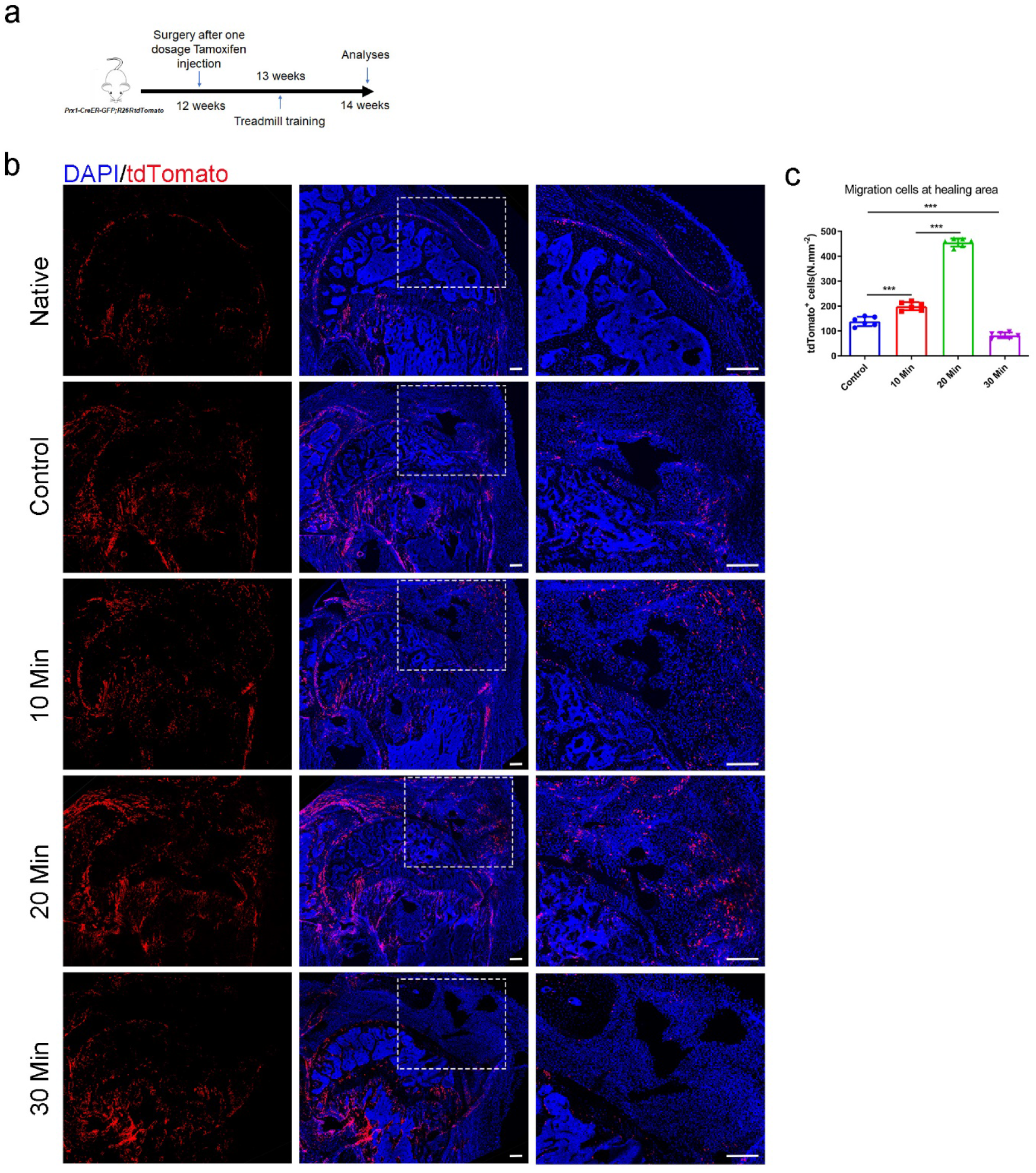
Proper mechanical stimuli could enhance the migration of Prx1^+^ cells to participate in enthesis healing. (a) Schematic diagram of *Prx1CreER-GFP; R26R-tdTomato* mice which were received enthesis surgery and sacrificed for immunofluorescent analysis at 14 weeks after surgery after sigle dose tamoxifen injection. (b) Representative immunofluorescent images of Tdtomato (red) staining of Prx1^+^ cells and DAPI (blue) staining of nuclei under different mechanical stimuli. Scale bar, 200 μm. (c) Quantitative analysis of migration Prx1^+^ cells at the healing area. n=6 per group. Scale bar, 200 μm. *P < 0.05, **P < 0.01, ***P < 0.001.

### TGF-β1 mediated mechanical stimulation to enhance enthesis injury repair

Previous report showed that TGF-β1 can recruit mesenchymal stem cells to maintain the balance of bone resorption and formation.^[37]^ The GO analysis found that Prx1^+^ cells were highly responded to TGF-β at P7, when enthesis initial mineralization began. Therefore, we hypothesized that TGF-β1 played an important role in enthesis repair process. First, we harvested the enthesis samples to perform ELISA analysis to reveal the content of active TGF-β1 during enthesis repair procedure (Figure 6a). We found that active TGF-β1 concentration increased and reached the peak at 2 weeks after surgery. Then, it returned to its basal level at 10 weeks. Meantime, mechanical stimulation could stimulate the release of active TGF-β1 during the repair procedure (Figure 6b).

**Figure 6.**
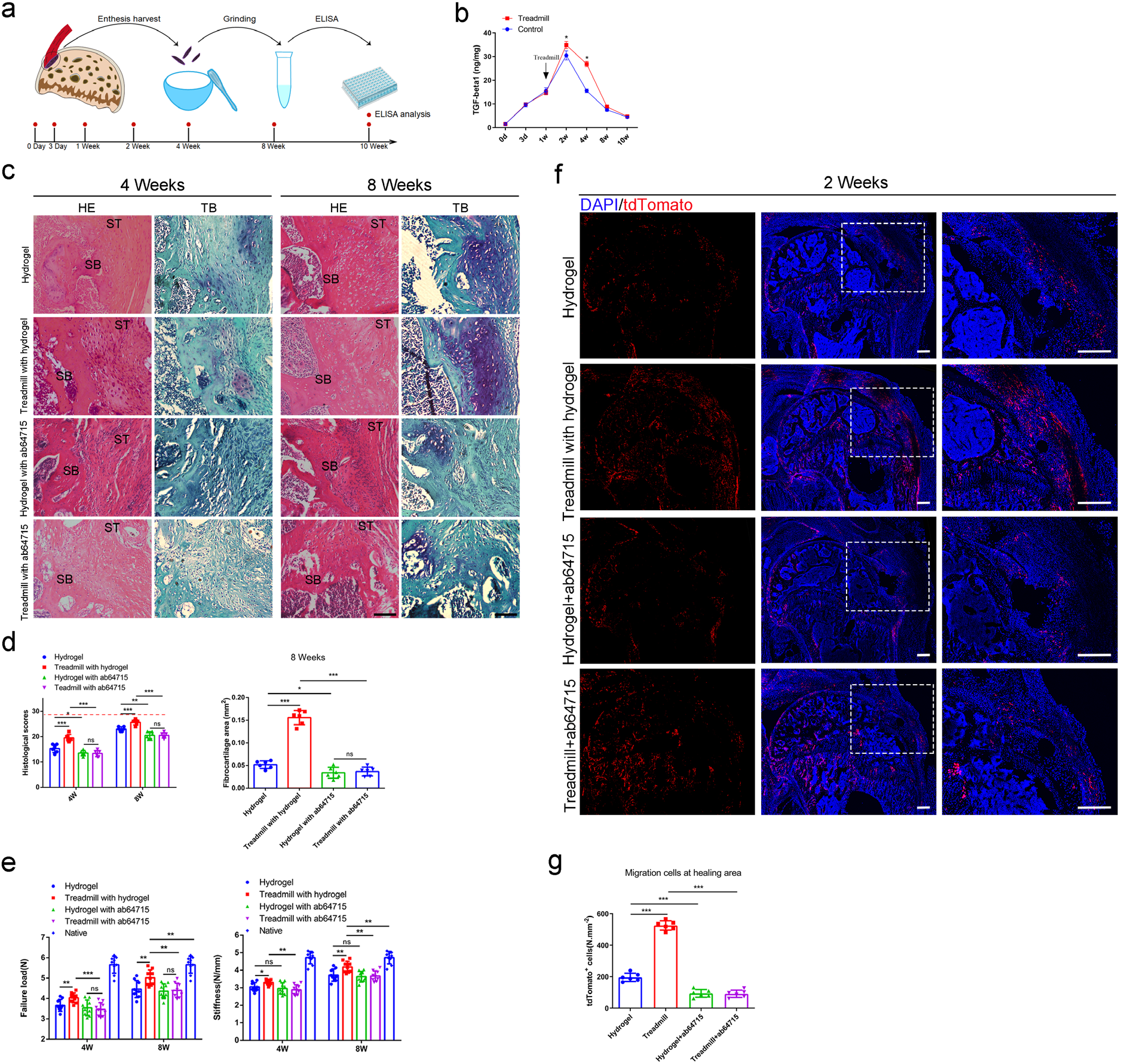
TGF-β1 mediated mechanical stimulation to enhance enthesis injury repair. (a) Schematic diagram of ELISA analysis. (b) ELISA analysis of TGF-β1 concentration during the enthesis healing procedure. n=3 per group. (c) Representative image of H&E and Toluidine blue/Fast green staining of enthesis. Scale bar, 200 μm. (d) Quantitative analysis of H&E score and fibrocartilage thickness. The red dolt line indicated the perfect histological score of 28. n=6 per group. (e) Quantitative analysis of Failure Load and stiffness. n=9 per group. (f) Representative immunofluorescent images of Tdtomato (red) staining of Prx1^+^ cells and DAPI (blue) staining of nuclei under different mechanical stimuli at postoperative 2 weeks. Scale bar, 200 μm. (g) Quantitative analysis of migration Prx1^+^ cells at the healing area. n=6 per group. *P < 0.05, **P < 0.01, ***P < 0.001.

To investigate the role of TGF-β1 during enthesis repair procedure, mice were recieved enthesis surgery and treated with or without TGF-β1 neutralizing antibody (ab64715). At 4 and 8 weeks, mice were sacrificed for histological and mechanical test analysis. The results showed that highly cellular, fibrovascular granulation tissue were observed at the supraspinatus enthesis in all groups at 4 weeks after surgery. Fibrovascular scar in the groups without ab64715 were relatively organized in H&E staining. Results of toluidine blue/fast green staining exhibited that mature fibrocartilage occurred more in groups without ab64715 than other groups with ab64715 (P<0.05 for all). Well-organized soft tissue and tidemark at the enthesis occurred at 8 weeks after surgery. However, enthesis in groups treated with ab64715 showed weak remodeling tissue than the groups without ab64715. The fibrocartilage was thinner in the groups with ab64715 than that in other groups without ab64715 (P<0.05 for all) (Figure 6c, d). The mechanical test showed that groups with ab64715 had lower failure load and stiffness than other groups without ab64715 at each time point (P<0.05 for all) (Figure 6e).

To understand if TGF-β1 also mediated the migration of Prx1^+^ cells to participate in enthesis injury repair, we performed lineage tracing analysis using *Prx1CreER*; *R26R-tdTomato* mice. After a single dose of tamoxifen (100 mg/kg, i.p.) injection, the mice were received surgery to create a enthesis repair model with or without ab64715 treatment. Mice began to receive treadmill training (20 minutes per day, five consecutive days per week) at 1 week after surgery and were sacrificed for immunofluorescence analysis at 2 weeks. We found that treadmill training could enhance Prx1^+^ cells to the healing area and this effect could be eliminated by the treatment with TGF-β1 neutralizing antibody (Figure 6f).

To find out if mechanical stimulation could have indipendent effect on Prx1^+^ cells migration, we isolated Prx1^+^ cells and investigated the effect of mechanical stimulation on Prx1^+^ cells with or without TGF-β1. We used a special dish that could load tensile force to Prx1^+^ cells (Figure S3a). After 4 consecutive days of mechanical stimuli (5%, 0.5 Hz, 20 minutes per day) with TGF-β1 (0.1 ng/ml), Prx1^+^ cell migration ability was analyzed by scratch assay and Transwell assay. We found that mechanical stimulation could not indipendently enhance the Prx1^+^ cell migration ability. At the same time, TGF-β1 could improve its migration ability, and this effect could be significantly stimulated by mechanical stimulation (Figure S3b, c, d). Western blot showed that pSmad2/3 was activated during this process (Figure S3e, f). These results indicated that treadmill training mobilised Prx1^+^ cells to enhance enthesis injury repair mainly by mediating the release of active TGF-β1.

### Primary cilia was essential for TGF-β signaling to promote enthesis injury repair

Previous studies showed that there were many receptors in primary cilia, which played an essential role in signal transmission.^[27,38–40]^ To determine whether the primary cilia plays an essential role in the transmission of TGF-β signaling, we created the primary cilia conditional knocked out transgenic mice. *Prx1CreER*; *IFT88*^*flox/flox*^; *R26R-tdTomato* mice and *Prx1CreER*; *R26R-tdTomato* mice were received enthesis injury repair surgery after 5 days continuous tamoxifen injection (75mg/Kg, i.p.). The mice were recieved treadmill training at the day 7 after surgery (20 minitues per day, 5 days per week) and sacrificed for assessment at 4 and 8 weeks. Results showed that conditional ablation of *IFT88* in Prx1^+^ cells significantly damaged the primary cilia (Figure 8a, b). Without the primary cilia, mechanical stimulation could not enhance the migration of Prx1^+^ cells to the healing area (Figure 8c, d). Results of H&E staining showed that less scar tissue formed at the enthesis at 4 weeks after surgery, and there was no significant difference in histological scores between the primary cilia dysfunction groups with or without treadmill training. No fibrocartilage was found in both groups at this time point. At 8 weeks after surgery, H&E staining showed high cellular, fibrovascular granulation tissue at the enthesis. Meanwhile, few fibrocartilage tissues were found at the enthesis site in these two *IFT88* damaged groups. There was no significant difference in H&E scores and the fibrocartilage area between the mice with or without treadmill training (Figure 8e, f, g). No significant difference in failure load and stiffness was found between the mice with or without treadmill training (Figure 8j).

### TGF-β1 enhanced the migration of Prx1^+^ cells via ciliary TGF-β signaling

To investigate the relationship between primary cilia and TGF-β signaling pathway, we isolated the Prx1^+^ cells and examined the distribution of TGF-β receptor 2 (TGF-βR2) in the cells with or without mechanical stimulation. Results showed that TGF-βR2 existed on the surface of Prx1^+^ cells. At the same time, TGF-βR2 was concentrated in the primary cilia under the effect of TGF-β1 (0.1 ng/ml), and mechanical stimuli could improve TGF-βR2 translocated into the primary cilia (Figure 7a, b). To understand if ciliary TGF-βR2 was essential for TGF-β signaling transmission, we used shRNA to knock down pallidin (PLDN) in Prx1^+^ cells, which could inhibit TGF-βR2 translocating into the primary cilia.^[28]^ Results showed that PLDN in Prx1^+^ cells could significantly be knocked down by shRNA (Figure 7c, d). TGF-βR2 concentrating in primary cilia was decreasing markedly in PLDN knocked down group (Figure 7e, f). The results of scratch assay and transwell assay showed that the effect of TGF-β1 on Prx1^+^ cells migration ability was eliminated in PLDN knocked down group (Figure 7g, h, i). Western blot analysis showed that the Smad2/3 signaling pathway was also inhibited at the same time (Figure 7j, k).

**Figure 7.**
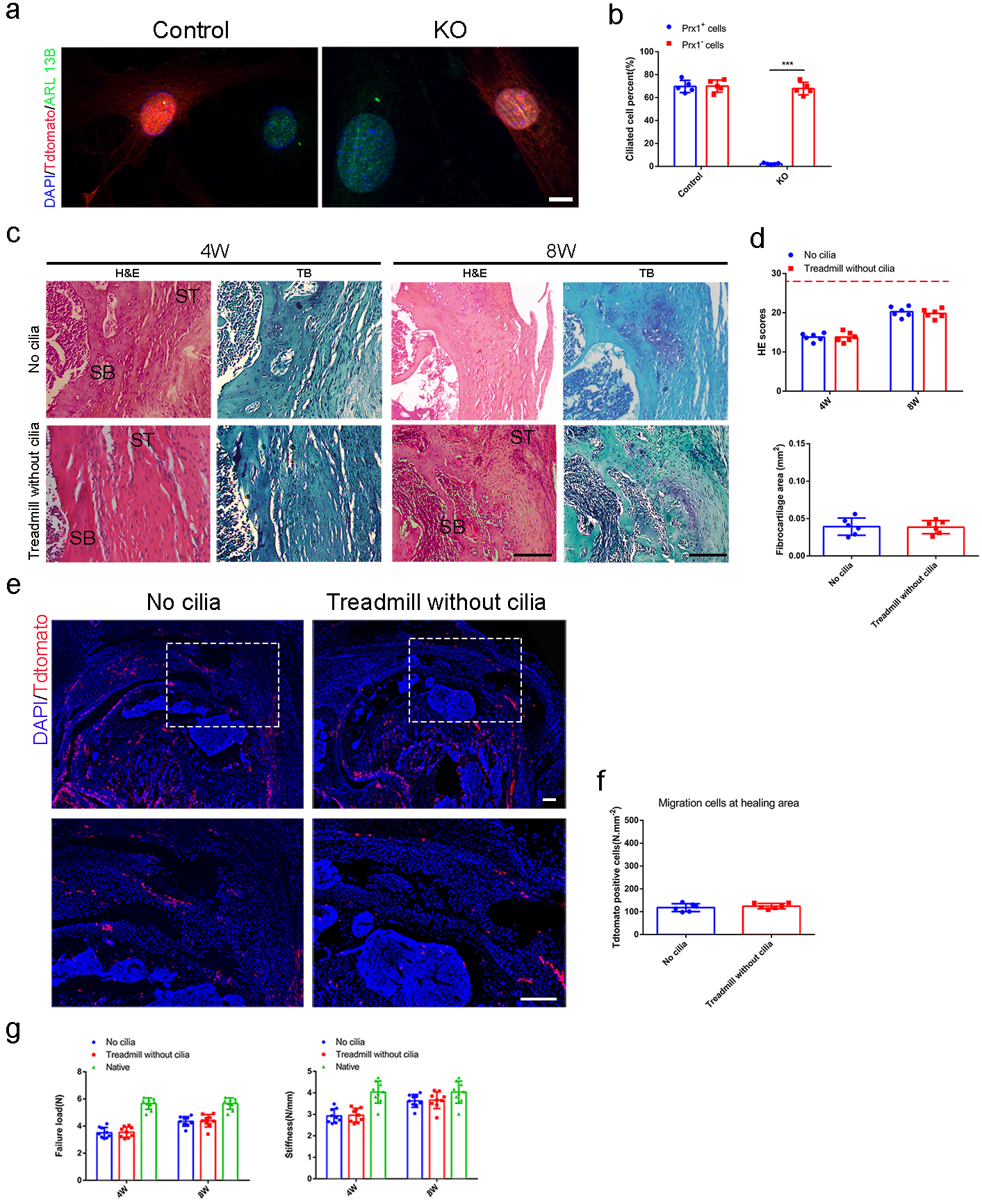
Primary cilia was essential for TGF-β signaling to promote enthesis injury repair. (a) Representative immunofluorescence image of tdTamato (red) staining of Prx1^+^ cells, ARL 13B (green) staining of primary cilia, and DAPI (blue) staining of nuclei. Scale bar, 5 μm. (b) Quantitative analyses of ciliated cell percent in Prx1^+^ cells and Prx1^−^ cells. n=5 per group. (c) Representative image of H&E and Toluidine blue/Fast green staining of enthesis. Scale bar, 200 μm. (d) Quantitative analysis of H&E score and fibrocartilage area at the enthesis. The red dotted line stands for perfect H&E scores of 28. n=6 per group. (e) Representative immunofluorescence image of tdTamato (red) staining of Prx1^+^ cells, DAPI (blue) staining of nuclei at the enthesis. Scale bar, 200 μm. (f) Quantitative analyses of Tdtomato^+^ cells in the healing area. n=6 per group. (g) Quantitative analyses of load failure and stiffness of enthesis. n=9 per group. SB, subchondral bone; ST: supraspinatus tendon. ***P < 0.001.

**Figure 8.**
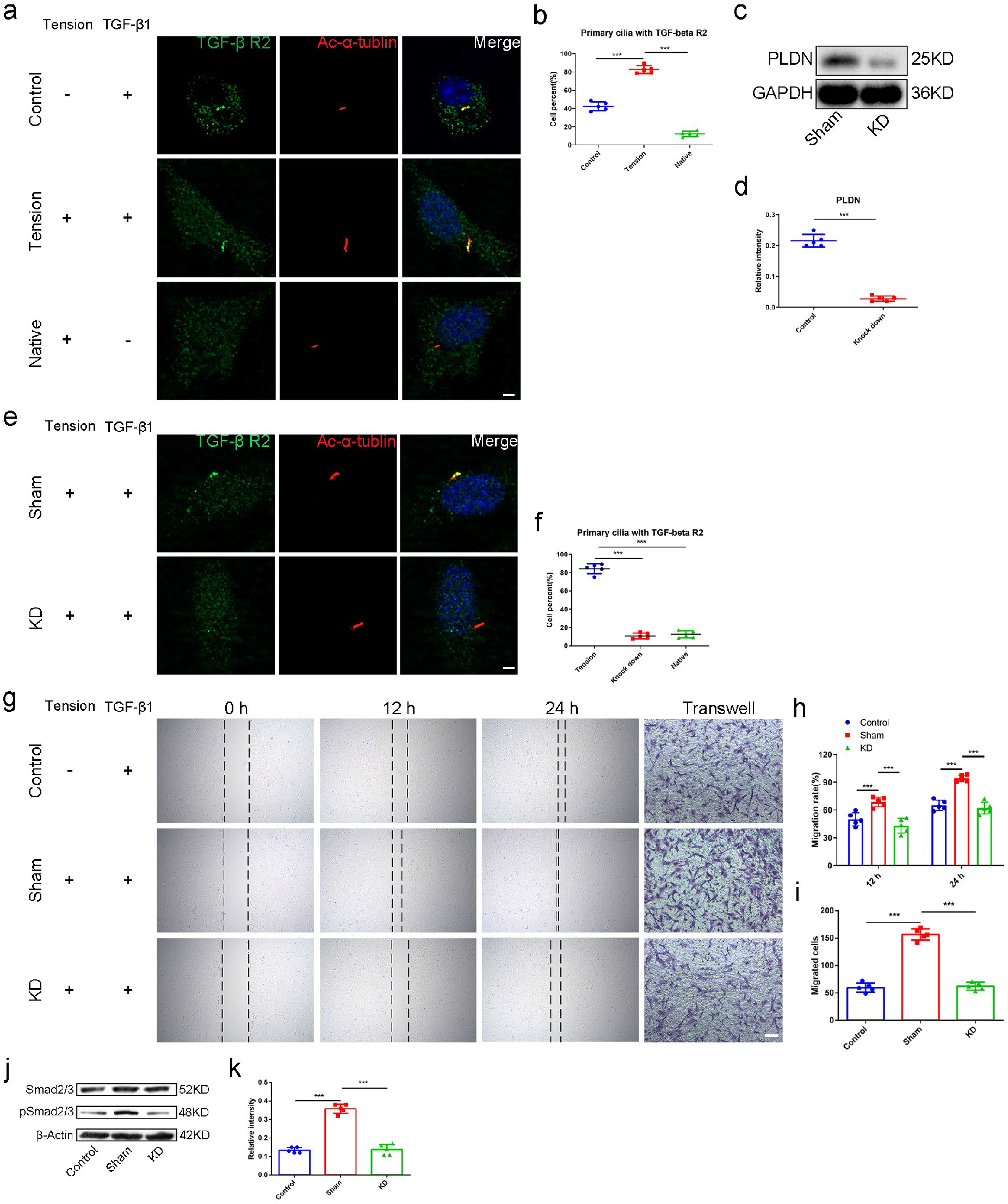
TGF-β1 enhanced the migration of Prx1^+^ cells via ciliary TGF-β signaling. (a) Representative image of immunofluorescent analysis of TGF-βR2 (green), Ac-α-tubulin (red) staining of primary cilia, and DAPI (blue) staining of nuclei stimulation by mechanical force with TGF-β1 (0.1 ng/ml) in Prx1^+^ cells. Scale bar, 5 μm. n=5 per group. (b) Quantitative analysis of cell percent that the TGF-βR2 was concentrated in the primary cilia. n=5 per group. (c) Western blot analysis of PLDN in groups treated with or without PLDN-siRNA. (d) Quantitative analysis of western blot. n=5 per group. (e) Representative image of immunofluorescent analysis of TGF-βR2 (green), Ac-α-tubulin (red) staining, and DAPI (blue) staining of nuclei stimulation by mechanical force with TGF-β1 (0.1 ng/ml) in Prx1^+^ cells treated with or without PLDN-siRNA. Scale bar, 5μm. (f) Quantitative analysis of cell percent that the TGF-βR2 was concentrated in the primary cilia. n=5 per group. (g) Scratch assay and transwell assay of Prx1^+^ cells. (h) Quantitative analyses of scratch assay. n=5 per group. (i) Quantitative analyses of transwell assay. n=5 per group. (j) Western blot analysis of Smad2/3/pSmad2/3 signaling. (k) Quantitative analyses of western blot. n=5 per group. *P < 0.05, **P < 0.01, ***P < 0.001.

## Discussion

Prx1^+^ cells and mechanical stimulation have been extensively studied during the skeletal development.^[41,42]^ However, their role in enthesis regeneration is poorly understood. In this study, we identified that Prx1^+^ cells are involved in enthesis development, but may not be important player in adult enthesis. However, during injury repair, Prx1^+^ cells could migrate to the injury area to participate in enthesis healing. Proper mechanical stimulation could increase the release of TGF-β1 to immobilize Prx1^+^ cells and promote enthesis injury repair. Ciliary TGF-βR2 was essential for TGF-β signaling transmission during proper mechanical stimulation promoting enthesis injury repair procedure. As far as we known, this is the first report uncovering the characteristics of Prx1^+^ cells on enthesis development and injury healing, and provided new insights into the progenitor source for enthesis injury repair. Meanwhile, this study found a new mechanism about the mechanical stimulation signal transmission, and had pieced together a mechanical conduction mechanism.

The microstructure regeneration of enthesis is a difficult issue considering the complicated structure of enthesis consisting of four continuous gradient layers: bone, calcified fibrocartilage, uncalcified fibrocartilage and tendon, and low self-regeneration ability.^[8]^ Therefore, current treatments aim to regenerate enthesis microstructure to acquire reliable long-term clinical results. A previous study found that low-intensity pulsed ultrasound stimulation after autologous adipose-derived stromal cell transplantation could improve the fibrocartilage and bone regeneration, leading to a better enthesis healing quality.^[43]^ Recently, tissue engineering was prevalent for repairing enthesis injury and showed promising results of fibrocartilage and bone regeneration, associated with better mechanical testing results.^[8,44,45]^ In the present studies, we found that better regeneration of fibrocartilage was higly correlated with the number of Prx1^+^ cells and was accompanied by better mechanical testing results, which was consistent with previous reports.These data suggested that the fibrocartilage regeneration directly influenced the enthesis healing quality and could be a reliable indicator for the enthesis regeneration.

Mechanical stimulation is a common therapy in the clinic, while it is a double-edged sword for enthesis healing. On one hand, appropriate mechanical load could stimulate local blood circulation, promote mesenchymal stem cell differentiation, and help releasing anti-inflammatory factors to improve tissue healing.^[46–48]^ Wang et al^[49]^ reported if the training started early, such as 96 hours after acute patellar tendon enthesis injury, it will lead to better recovery results with fibrocartilage and mechanical test parameters in a rabbit model. On the other hand, excessive mechanical loading in the early healing phase might delay or even inhibit tissue healing.^[14]^ Rodeo et al found that immediate excessive treadmill training at the early stage causes delayed enthesis healing in the murine enthesis repair model.^[50]^ These reports suggested that enthesis could only respond favorably to controlled loading after injury. In the present studies, we chose a 7-days delayed mechanical stimulation and tested 3 sets of mechanical intensity. We found that 20 minutes per day mechanical stimulation (7-days delayed, 10 m/min, 5 days per week) could enhance Prx1^+^ cell migration, improve the quality of fibrocartilage and bone regeneration in the enthesis, resulting in better biomechanical results. Our finding suggested that appropriate mechanical stimulation (7-days delayed, 10 m/min, 20 minutes per day, five days per week) indeed improve early enthesis healing, and this model was reliable to uncover the mechanism of mechanical stimulation-induced enthesis healing.

Our results showed that proper mechanical stimuli could enhance the migration of Prx1^+^ cells and Prx1^+^ cells could differentiate in cartilage and bone cells which play a critical role in enthesis healing. We know that Prx1^+^ cells are important in skeletal development, especially during the embryonic development.^[17]^ However, their role in enthesis development and contribution to skeletal tissue regeneration was poorly understood. In this study, we found that GFP^+^ cells (active Prx1^+^ cells) were abundant in the enthesis during early stage and disappeared at adulthood. The tdtomato^+^ cells mainly localized within the periosteum, perichondrium, and growth plate at adulthood, which indicated that Prx1^+^ cells confined at these places were active in young age and quiescence, but not completely disappear at adulthood. Meanwhile, there were no Prx1^+^ cells at the enthesis area in adult mice. When enthesis injury happens, Prx1^+^ cells could migrate to the injury area to participate in the healing process. Besides, our results showed that Prx1^+^ cell numbers at the enthesis were related to the healing quality, suggesting that Prx1^+^ cells were pivotal for enthesis healing. One limitation of this study is that we did not investigate other mesenchymal sub-population in the current study.

How did mechanical stimulation enhance the migration of Prx1^+^ cells to the healing site? As we known, TGF-β signaling plays an essential role in tissue homeostasis and injury healing and TGF-β signaling activation is in precise spatial and temporal manner.^[51–53]^ The level of TGF-β1 was relatively high at the healing interface and TGF-β1 could promote the migration of MSCs to modulate bone remodeling.^[37]^ Robertson et al reported that TGF-β1 function is mainly regulated by its activation rather than synthesis or secretion.^[54]^ Hence, we mainly focused on the activation of TGF-β1. Our results showed that TGF-β signaling is involved in enthesis healing. During this process, active TGF-β1 concentration was elevated, and was further enhanced by mechanical stimulation. In vitro, we found that mechanicla stimulation couldn’t indipendently improve Prx1^+^ cells migration ability, while it could enhance the sensitivity of the Prx1^+^ cells to TGF-β1. Although we didn’t further reveal its effects on Prx1^+^ cells migration in vivo, it still provided new insights in new mechanisms about the mechanical stimulation signal transmission. At the same time, we could not rule out the involvement of other signaling pathways in this process.

Primary cilia have been recognized as an essential cellular mechanoreceptor and mechanosensitive channels.^[55,56]^ Still, the regulatory mechanism of primary cilia in mechanical stimulation transmission remains unclear, even controversial. Polycystin-1 (PC1) and polycystin-2 (PC2), co-distribution in the primary cilia of kidney epithelium, was reported to be involved in the regulation of intracellular Ca^2+^ signaling and transfer mechanical stimuli.^[24]^ In addition to PC1 and PC2, another primary cilia-based calcium channels-transient receptor potential (TRP) could also sense mechanical stimuli via conducting Ca^2+^ signaling.^[57]^ Nonetheless, many calcium channels are not only localized in the ciliary membrane, but also other parts of the cells, so it is difficult to differentiate the difference between ciliary and cytosolic Ca^2+^ in response to the same mechanical stimuli. Delling et al found that cilia-specific Ca^2+^ influxes were not observed in physiological or even highly supraphysiological levels of fluid flow.^[58]^ In this study, we found that knocking out IFT88 prevented the mechanotransduction. Inhibition of ciliary TGF-β signaling could decrease the mechanotransduction, suggesting that primary cilia could regulate mechanical stimuli via ciliary TGF-βR2. This finding provides new insights into the role of primary cilia in mechanical stimuli transmission. However, we didn’t exclude ciliary Ca^2+^ signaling or other ciliary signaling pathway participating in this mechanical stimulation transmission process in this study.

## Conclusion

In conclusion, Prx1^+^ cells were an essential subpopulation of progenitors for enthesis development and injury repair. Mechanical stimulation could increase the release of TGF-β1 and enhance mobilization of Prx1^+^ cells to promote enthesis injury repair via ciliary TGF-β signaling.

## Experimental Section/Methods

### Collection of Single-Cell Suspension from Supraspinatus enthesis

In general, 10-12 suspensions tendon enthesis tissue (E15.5, P7 and P28), including the tendon (one millimeter in length) and the portion of the humeral head proximal to the growth plate near the tendon attachment, were collected from pooled sibling shoulders (five to six mice per pool). Fresh enthesis tissue were finely chopped with small scissors in 1 ml of Dulbecco’s modified Eagle’s medium (DMEM), then digested in 0.5 % type I collagenase (Life Technologies) and 7 U/ml Dispase II (Gibco) at 37 °C for 30 min. Then the supernatant was collected and filtered through 70 μm cell filters (Falcon BD), and centrifuged for 5 min at 300 g, before re-suspending the pellet in DMEM containing 2 % serum, and the cell suspension was kept on ice until load on chip.

### Flow Cytometry and Cell Sorting

Collected cell suspension were blocked with purified rat anti-mouse CD16/CD32 (BD Pharmingen, dilution 1:100) for 10 min, then stained with fluorophore conjugated antibodies. Antibodies used in this study are anti-mouse CD45-APCCy7, Ter119-APCCy7 (Biolengend), DAPI (eBiosciences) stain was used to exclude dead cells. For cell sorting, single cells were gated using doublet-discrimination parameters and cells were collected in FACS buffer (1x HBSS, 2 % FBS, 1 mM EDTA). Cell viability was assessed with trypan blue and only samples with > 85% viability were processed for further sequencing.

### scRNA-Seq Sequencing, Data processing and quality control

Around 10,000 sorted live CD45-Ter119- cells for each timepoint sample were resuspended in FACS buffer according to the recommendations provide by 10× Genomics for optimal cell recovery. Single-cell mRNA libraries were built using the Chromium Single Cell 3’ kit (v3), libraries sequenced on an Illumina NovaSeq 500 instrument. Single-cell fastq sequencing reads from each sample were processed by aligning reads and obtaining unique molecular identifier (UMI) counts and converted to gene expression matrices, after mapping to the mouse (mm10) reference genome using the Cell Ranger v4.0.0 pipeline, according to the standard workflow (10× Genomics).

### scRNA-seq Data Ananlysis

Quality control was conducted for each dataset, cells with less than 200 genes and the top 10% cells were removed to minimize multiplet possibility. Cells were retained if the percent mitochondrial reads were lower that 20% (8919 cells for embryonic day 15.5, 7489 cells for postnatal day 7 and 5124 cells for postnatal day 28). Data integration, graph-based cell clustering, dimensionality reduction, and data visualization were analyzed by the Seurat R package (v3.2). Data integration was performed via canonical correlation analysis to remove batch effect. Feature (gene) data was scaled in order to remove unwanted sources of variation using the Seurat ScaleData function for percent mitochondrial reads, number of genes detected and predicted cell cycle phase difference. Non-linear dimension reduction was performed using uniform manifold projection (UMAP) and graph-based clustering was performed using the Louvain algorithm. The number of statistically significant principal components were set empirically by testing top 10 differentially expressed genes (MAST method) between the clusters. Subsetting was performed by assessing marker gene expression across clusters, Clusters associated with the following cell-types were excluded from our analysis: muscle cells, immune cells blood cells, endothelial cells and undefined cells rich of histone genes. Functional annotation of the marker genes relative to GO terms was performed using ClusterProfiler (v3.18). Trajectory analysis of the tendon enthesis development was performed using Monocle (v2.4.0).

### Animals and treatment

The *Prx1CreER-GFP* (Strain origin: C57BL6N/129; Stock No: 029211), *IFT88*^*flox/flox*^ (Strain origin: C57BL6N/129; Stock No: 022409); *Rosa26tdTomato* (Strain origin: C57BL6N/129; Stock No: 007909) mouse strain was purchased from Jackson Laboratory (Bar Harbor, ME).

*Prx1CreER-GFP* mice were crossed with *IFT88*^*flox/flox*^ mice. The offspring were intercrossed to generate the following genotypes: WT, *Prx1CreER*-GFP, *Prx1CreER-GFP*; *ITF88*^*flox/flox*^. Then, *Prx1CreER-GFP*; *ITF88*^*flox/flox*^ mice were crossed with *Rosa26tdTomato* mice. The offspring were intercrossed to generate the following genotypes: *Prx1CreER-GFP*; *Rosa26tdTomato* mice (mice expressing tdTomato driven by Cre recombinase in Prx1^+^ cells), *Prx1CreER-GFP*; *ITF88*^*flox/flox*^; *Rosa26tdTomato* mice (conditional deletion of *ITF88* in Prx1 lineage cells and expressing tdTomato, referred to as *IFT88*^−/−^ herein). To induce Cre recombinase activity, we injected mice at designated time points with tamoxifen (75 mg/kg, i.p.) for consecutive five days.

The genotype of the mice was determined by PCR analysis of genomic DNA, extracted from mouse tails using the primers as follows. *Prx1CreER-GFP* allele forward, 5′-ATACCGGAGATCATGCAAGC-3′, reverse, 5′-GGCCAGGCTGTTCTT CTTAG-3′, control forward, 5′-CTAGGCCACAGAATTGAAAGATCT-3′ and control reverse, 5′-GTAGGTGGAAATTCTAGCATCATCC-3′; *IFT88*^−/−^ allele forward, 5′-TGAGGACGACCTTTACTCTGG-3’, and reverse, 5′-CTGCCATGACTGGTTCT CACT-3′; *Rosa26tdTomato* allele forward, 5′-AAGGGAGCTGCAGTGGAGTA-3′, reverse, 5′-CCGAAAATCTGTGGGAAGTC-3′, control forward, 5′-GGCATTAAAG CAGCGTATCC-3′ and control reverse, 5′-CTGTTCCTGTACGGCATGG-3′.

### Rotator cuff injury repair model

Twelve weeks old male mice underwent rotator cuff injury repair using protocol as previously reported.^[15,59]^ After anesthetized with pentobarbital (0.6 mL/20 g; Sigma-Aldrich, St. Louis, MO), a longitudinal skin incision was made to expose the deltoid muscle, and a transverse cut was made on it. The acromion was pulled away to expose the supraspinatus tendon. After the supraspinatus tendon was grasped with 6-0 Prolene (Ethicon, Somerville, NJ, USA), it was sharply transected at the insertion site on the greater tuberosity, and fibrocartilage layer was removed with a scalpel blade. A bone tunnel was made transversely to the distal greater tuberosity. Then, the suture was passed through the drilled hole and tied the supraspinatus tendon to its anatomic position. The skin and deltoid muscles were closed in layer. To block the TGF-β1, hydrogel loading with the TGF-β1 neutralizing antibody was used. Mice were allowed free cage activities. At postoperative four weeks and eight weeks, the mice receiving treadmill exercise or not were sacrificed, and the supraspinatus-humeral head composites were harvested for further study.

### Mechanical load in vivo and in vitro

A motor-powered treadmill with 12 lanes was used to generate a mechanical load to the enthesis in vivo. Treadmill exercise was conducted as previously reported.^[15,50]^ All the mice underwent one-week adaptive training to get familiar with the lane environment before surgery. Treadmill speed was increased daily until all mice tolerated running at 10 m/min for 30 minutes per day. At postoperative day 7, the mice in the treadmill group ran at a speed of 10 m/min on a 0° declined lane for 10 minutes, 20 minutes or 30 minutes, five days per week.

A cell load system (CellLoad-300, Hao Mian, China) was used to generate tensile mechanical load on Prx1^+^ cells. Prx1^+^ cells were seeded on a plate which could expand and contract under external forces, at a density of 1.5×10^4^/cm^2^. The parameters were set as follows: 1Hz, 5%, 20 minutes per day.

### Immunofluorescence

Humeral head and supraspinatus tendon composite were harvested and fixed in the 4% paraformaldehyde in PBS overnight at room temperature. After decalcified and dehydrated, samples were embedded in Tissue-Tek^®^ O.C.T. Compound (SAKURA, Torrance, USA) and cut into 10 μm thickness of sagittal sections. Cell samples were fixed in the 4% paraformaldehyde in PBS for 30 minutes at room temperature. Both the parts and cell samples were blocked in 5% BSA for 40 min at room temperature and incubated with the primary antibodies anti-DMP1 (Abcam, 1:400, ab13970), anti-GFP (Abcam, 1:400, ab13970 or 1:400, ab290), anti-Sox9 (Abcam, 1:400; ab185966), anti-TGF-βR2 (Abcam, 1:400, ab186838) at 4°C overnight. After washing, the sections were then incubated with the respective secondary antibodies (1:500, Abcam) for 1 hour at room temperature and sealed with DAPI. The images were captured with a Leica TCS-SP8 confocal microscope (Leica, Germany).

### Histological analysis

After radiographic assay, fixed samples were decalcified in EDTA for 14 days, dehydrated in gradient ethanol, embedded in paraffin, and then cut into 5μm slices. The sections were stained with hematoxylin and eosin for general histology analyses. Two blinded observers measured histological tendon maturing score according to a previous report (Table S1)^[50]^.

### Biomechanical test

An Instron biomechanical testing system (Model 5942, Instron, MA) was used to detect the failure load and stiffness of these samples. The tendon was secured in a clamp using sandpaper, while the humerus firmly clamped with a vice grip. The specimens were tested at room temperature, and samples were preconditioned with 0.1 N and then loaded to failure at a rate of 0.1 mm/s. A consistent gauge length was used throughout the test. Data were excluded if the tendon slipped out of the grip or did not fail at the supraspinatus tendon attachment site.

### ELISA

The supraspinatus tendon insertion specimens were harvested at postoperative 0 day, and 3 days, and 1, 2, 4, 8, and 10 weeks. We removed the muscle belly and kept the tendon and the portion of the humeral head proximal to the growth plate near the tendon attachment. Then, we determine the concentration of active TGF-β1 in the enthesis using the ELISA Development Kit (R&D Systems, Minneapolis, MN) according to the manufacturer’s instructions.

### Cell culture

To obtain Prx1^+^ cells, 1-2 weeks old *Prx1CreER-GFP* mice were sacrificed. The tibiae and femurs were dissected and excised into chips of approximately 1-3 mm^3^ with scissors. Then, the chips were suspended into a 25 cm^2^ plastic culture flask with 5 ml of α-MEM containing 10% (vol/vol) FBS in the presence of 3 mg/ml (wt/vol) of collagenase II (Sigma) and digested the chips for one h in a shaking incubator at 37°C with a shaking speed of 150 rpm. Washed the enzyme-treated chips with α-MEM and got the Prx1^+^ cells by FACS. Reseeded the cells and changed the medium every 48h. In the same way, we isolated the Prx1^+^ cells without cilia through *Prx1CreER-GFP*; *IFT88*^*flox/flox*^ mice and *Prx1CreER-GFP*; *IFT88*^*flox/flox*^; *Rosa26tdTomoto* mice after tamoxifen injection for 5 days (75 mg/kg, i.p.).

### Short hairpin RNAs transfection

Prx1^+^ cells were transfected with shRNA targeting PLDN (shRNA#1: 5′-ATACACTGGAACAAGAGATTT-3’, shRNA#2: 5′-CGCCAAGCTGGTGACTAT AAG-3’) or with a scrambled shRNA for 12 hours using lentiviral vector (VectorBuilder, Cyagen Biosciences, Santa Clara, CA) at an MOI of 20. Prx1^+^ cells were maintained in growth media for a further 72 hours before the application of a mechanical stimulation.

### Scratch assay

For scratch wound assay, Prx1^+^ cells (1.5×10^4^ cells/cm^2^) were seeded into a stretchable dish and cultured with tension load (5%, 1Hz, 20 minutes per day) for three days. Cell monolayer could be formed at this time point. A straight scratch was produced using a pipette tip. After washed with PBS to remove floating cells, adherent, complete medium was added. Wound closure was imaged at the 0, 12, and 24 h of incubation time points. The rate of wound closure was calculated as follows: Migration rate (%) = (A0 – An)/A0 × 100, where A0 represents the initial wound area, and An represents the remaining area of the wound at the appointed time.

### Transwell assay

After tension force load for 4 days, 1×10^4^ Prx1^+^ cells were resuspended in 100μl α-MEM medium were loaded into the upper chamber of 24-well Transwell plate (Corning, NY, USA) with 8μm pore-sized filters. Complete medium, which supplemented with containing 0.1 ng/ml TGF-β1, was added to the lower chamber. After 12 h of incubation, cells that migrated to the lower surface of the filter were rinsed, fixed, and stained with 1% crystal violet. An optical microscope was used to photograph and count the migrated cells.

### Western blotting

Prx1^+^ cells with or without tension force load were collected for extracting protein. Western blotting was performed with 10% sodium dodecyl sulfate-polyacrylamide gel electrophoresis. Then the proteins were transferred into a nitrocellulose membrane, and membranes were blocked by nonfat milk. After blocking, the nitrocellulose membranes were then incubated using anti-PLDN (Proteintech, 1:500, 10891-2-AP), anti-Smad2/3 Ab (Abcam, 1:500, ab202445), anti-pSmad2/3 Ab (Abcam, 1:500, ab63399), anti-GAPDH (Proteintech, 1:1000, 110494-1-AP). The figures for western blotting were visualized using enhanced chemiluminescence reagent (Thermo Fisher Scientific, Waltham, USA) and imaged by the ChemiDoc XRS Plus luminescent image analyzer (Bio-Rad).

### Statistical analysis

The statistical results were analyzed by GraphPad Prism 7.0 software. Quantitative data were expressed as mean ± standard deviation (SD), and differences above 2 groups were evaluated using one-way ANOVA with post hoc test, while the histological scores was performed using the Mann-Whitney test. Statistical significance was set at *P*<0.05.

### Study approval

All animal care protocols and experiments in this study were reviewed and approved by the Animal Care and Use Committees of the Laboratory Animal Research Center of our institute. All mice were maintained in the specific pathogen-free facility of the Laboratory Animal Research Center.

## Acknowledgements

This work was supported by National Natural Science Foundation of China (No. 81730068) and the Major Science and Technology project of Changsha Science and Technology Bureau (NO.41965).. Thanks to professor Xianghang Luo and Hui Xie for supporting of this research.

## Supporting Information

**Figure S1.**
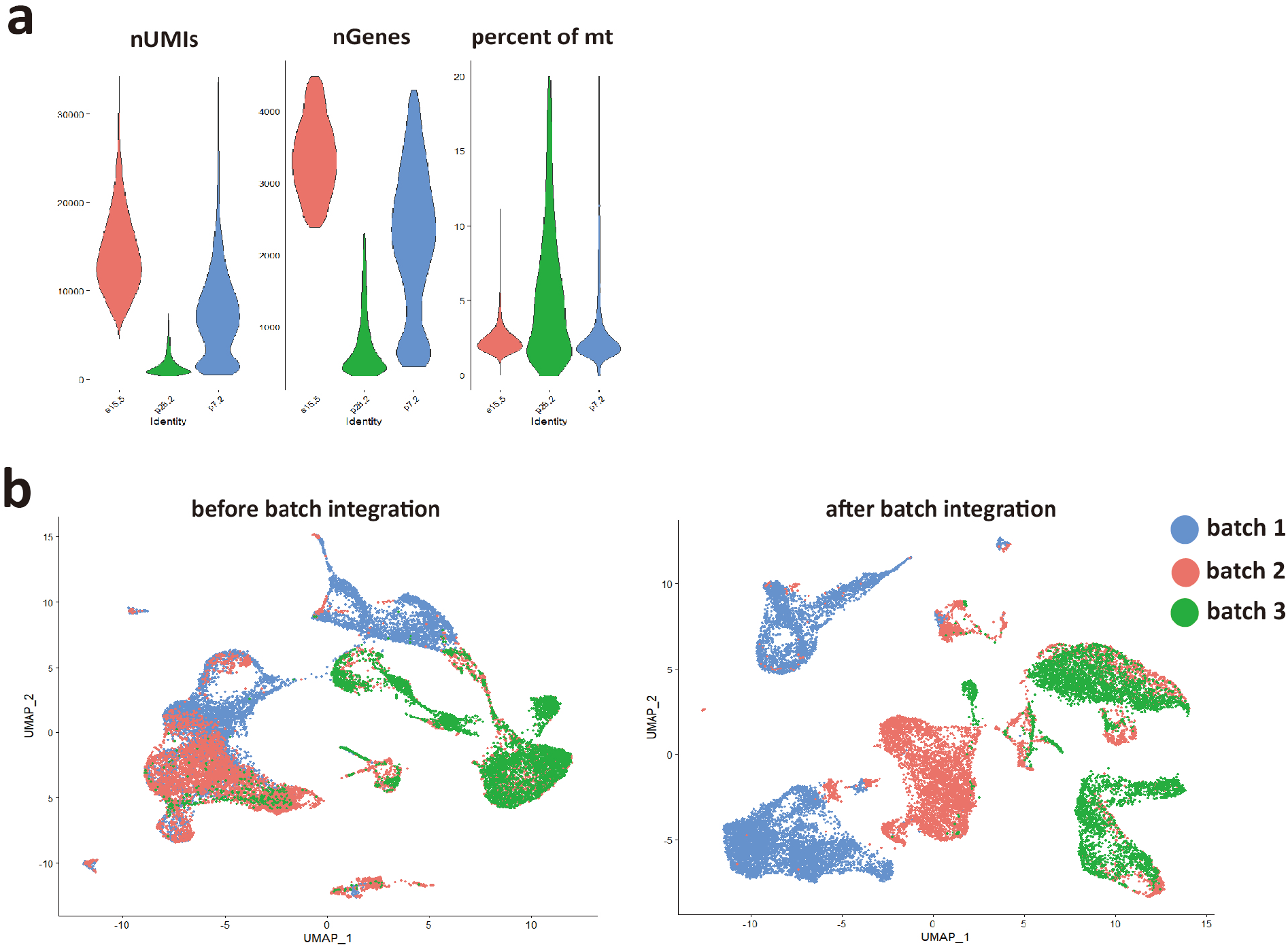
Quality control of unbiased scRNA-seq dataset. (a) Metrics used to assess the quality of the scRNA-seq libraries. Cells with less than 200 genes and the top 10% cells were removed to minimize multiplet possibility. (b) Batch effect was eliminated after data integration using Seurat.

**Figure S2.**
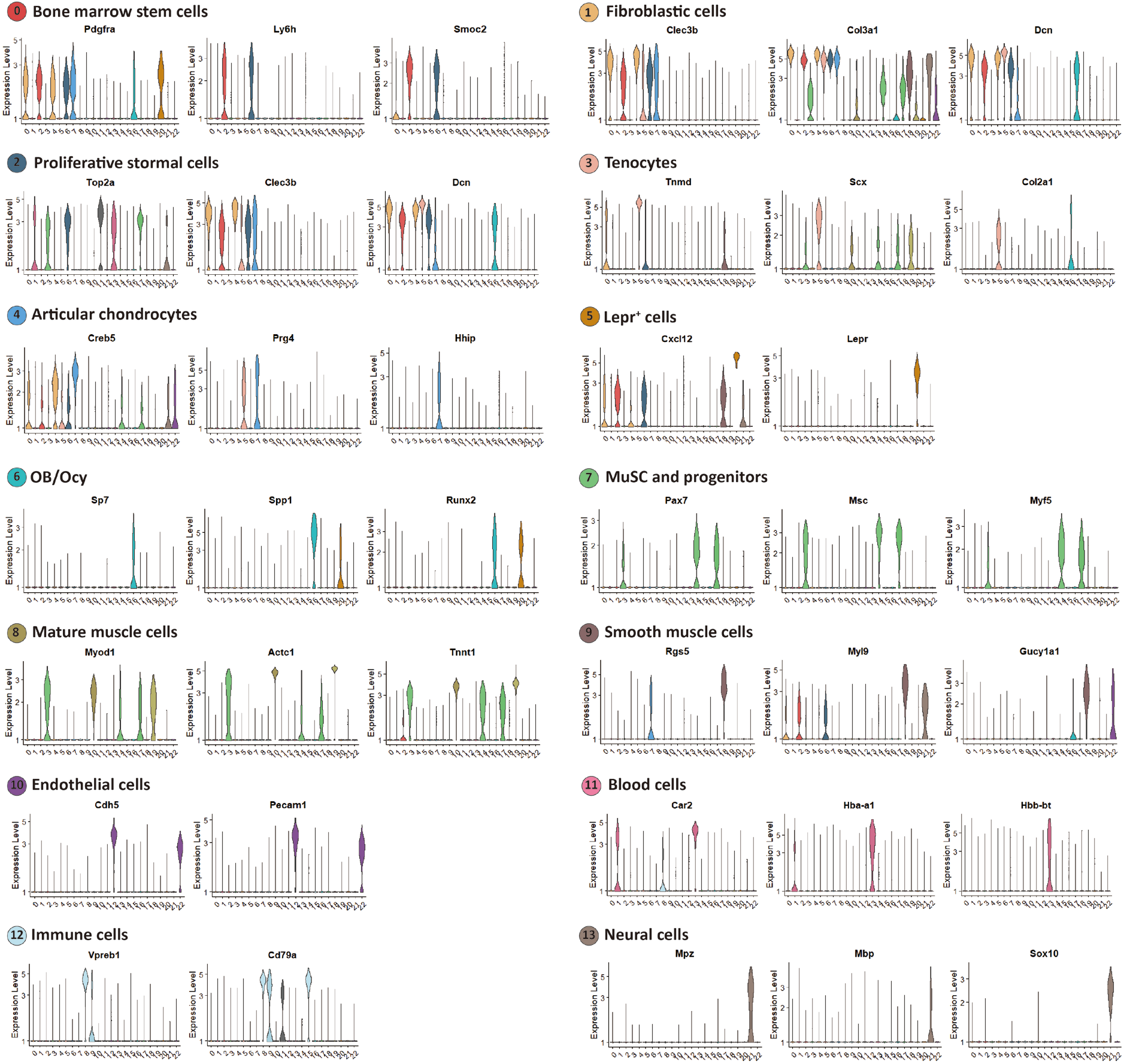
Gene expression defining clusters of E15.5, P7, P28 canonical correlation analysis.

**Figure S3.**
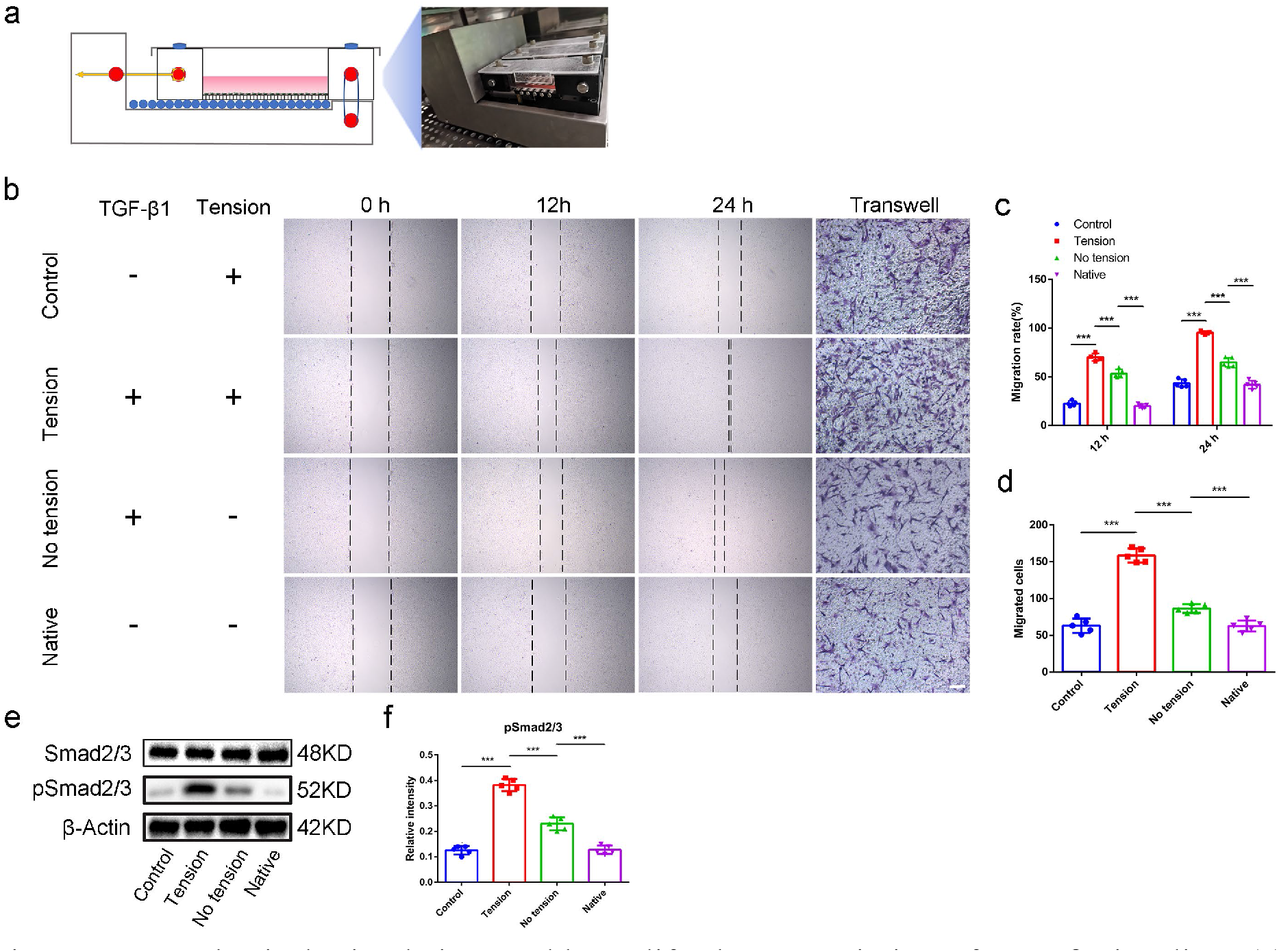
Mechanical stimulation could amplify the transmission of TGF-β signaling. (a) Schematic image and gross view of the cell loading system. (b) Scratch assay and transwell assay of Prx1+ cells. Scale bar, 100 μm. (c) Quantitative analyses of migration cells in a scratch assay. n=5 per group. (d) Quantitative analyses of migration cells in transwell assay. n=5 per group. (e) Western blot analyses of Smad2/3, phosphorylation of Smad2/3. (f) Quantitative analyses of western blot. n=5 per group. *P < 0.05, **P < 0.01, ***P < 0.001.

## REFERENCES AND NOTES

1. Meislin, R. J., Sperling, J. W. & Stitik, T. P. Persistent shoulder pain: epidemiology, pathophysiology, and diagnosis. Am J. Orthop. 34, 5–9 (2005).

2. Galatz, L. M., Griggs, S., Cameron, B. D. & Iannotti, J. P. Prospective longitudinal analysis of postoperative shoulder function : a ten-year follow-up study of full-thickness rotator cuff tears. J Bone Joint Surg Am 83, 1052–1056 (2001).

3. Chung, S. W., Kim, J. Y., Kim, M. H., Kim, S. H. & Oh, J. H. Arthroscopic repair of massive rotator cuff tears: outcome and analysis of factors associated with healing failure or poor postoperative function. Am J Sports Med 41, 1674–1683 (2013).

4. Thigpen, C. A., Shaffer, M. A., Gaunt, B. W., Leggin, B. G., Williams, G. R. & Wilcox, R. B. 3rd The American Society of Shoulder and Elbow Therapists’ consensus statement on rehabilitation following arthroscopic rotator cuff repair. J Shoulder Elbow Surg 25, 521–535 (2016).

5. Lee, S., Park, I., Lee, H. A. & Shin, S. J. Factors related to symptomatic failed rotator cuff repair leading to revision surgeries after primary arthroscopic surgery. Arthroscopy, (2020).

6. Hein, J., Reilly, J. M., Chae, J., Maerz, T. & Anderson, K. Retear Rates After Arthroscopic Single-Row, Double-Row, and Suture Bridge Rotator Cuff Repair at a Minimum of 1 Year of Imaging Follow-up: A Systematic Review. Arthroscopy 31, 2274–2281 (2015).

7. Galatz, L. M., Ball, C. M., Teefey, S. A., Middleton, W. D. & Yamaguchi, K. The outcome and repair integrity of completely arthroscopically repaired large and massive rotator cuff tears. J Bone Joint Surg Am 86, 219–224 (2004).

8. Chen, C. et al. Book-Shaped Acellular Fibrocartilage Scaffold with Cell-loading Capability and Chondrogenic Inducibility for Tissue-Engineered Fibrocartilage and Bone-Tendon Healing. ACS Appl Mater Interfaces 11, 2891–2907 (2019).

9. B. P. Thampatty, J. H. Wang, Mechanobiology of young and aging tendons: In vivo studies with treadmill running. J. Orthop. Res. 36, 557–565 (2018).

10. Thomopoulos, S., Kim, H. M., Rothermich, S. Y., Biederstadt, C., Das, R. & Galatz, L. M. Decreased muscle loading delays maturation of the tendon enthesis during postnatal development. J. Orthop. Res. 25, 1154–1163 (2007).

11. Galloway, M. T., Lalley, A. L. & Shearn, J. T. The role of mechanical loading in tendon development, maintenance, injury, and repair. J Bone Joint Surg Am 95, 1620–1628 (2013).

12. Chang, K. V., Hung, C. Y., Han, D. S., Chen, W. S., Wang, T. G. & Chien, K. L. Early Versus Delayed Passive Range of Motion Exercise for Arthroscopic Rotator Cuff Repair: A Meta-analysis of Randomized Controlled Trials. Am J Sports Med 43, 1265–1273 (2015).

13. Keener, J. D., Galatz, L. M., Stobbs-Cucchi, G., Patton, R. & Yamaguchi, K. Rehabilitation following arthroscopic rotator cuff repair: a prospective randomized trial of immobilization compared with early motion. J Bone Joint Surg Am 96, 11–19 (2014).

14. Lee, B. G., Cho, N. S. & Rhee, Y. G. Effect of two rehabilitation protocols on range of motion and healing rates after arthroscopic rotator cuff repair: aggressive versus limited early passive exercises. Arthroscopy 28, 34–42 (2012).

15. Zhang, T. et al. Treadmill exercise facilitated rotator cuff healing is coupled with regulating periphery neuropeptides expression in a murine model. J. Orthop. Res., (2020).

16. T. A. Wynn, K. M. Vannella, Macrophages in Tissue Repair, Regeneration, and Fibrosis. Immunity 44, 450–462 (2016).

17. Kawanami, A., Matsushita, T., Chan, Y. Y. & Murakami, S. Mice expressing GFP and CreER in osteochondro progenitor cells in the periosteum. Biochem. Biophys. Res. Commun. 386, 477–482 (2009).

18. Ouyang, Z. et al. Prx1 and 3.2kb Col1a1 promoters target distinct bone cell populations in transgenic mice. Bone 58, 136–145 (2014).

19. Martin, J. F. & Olson, E. N. Identification of a prx1 limb enhancer. Genesis 26, 225–229 (2000).

20. Nauli, S. M., Kawanabe, Y., Kaminski, J. J., Pearce, W. J., Ingber, D. E. & Zhou, J. Endothelial cilia are fluid shear sensors that regulate calcium signaling and nitric oxide production through polycystin-1. Circulation 117, 1161–1171 (2008).

21. Bowie, E. & Goetz, S. C. TTBK2 and primary cilia are essential for the connectivity and survival of cerebellar Purkinje neurons. Elife 9, (2020).

22. Hilgendorf, K. I. et al. Omega-3 Fatty Acids Activate Ciliary FFAR4 to Control Adipogenesis. Cell 179, 1289–1305.e21 (2019).

23. Vion, A. C. et al. Primary cilia sensitize endothelial cells to BMP and prevent excessive vascular regression. J. Cell Biol. 217, 1651–1665 (2018).

24. Nauli, S. M. et al. Polycystins 1 and 2 mediate mechanosensation in the primary cilium of kidney cells. Nat. Genet. 33, 129–137 (2003).

25. Hua, K. & Ferland, R. J. Primary cilia proteins: ciliary and extraciliary sites and functions. Cell. Mol. Life Sci. 75, 1521–1540 (2018).

26. Guemez-Gamboa, A., Coufal, N. G. & Gleeson, J. G. Primary cilia in the developing and mature brain. Neuron 82, 511–521 (2014).

27. Anvarian, Z., Mykytyn, K., Mukhopadhyay, S., Pedersen, L. B. & Christensen, S. T. Cellular signalling by primary cilia in development, organ function and disease. Nat Rev Nephrol 15, 199–219 (2019).

28. L. Zheng, Y. Cao, S. Ni, et al., Ciliary parathyroid hormone signaling activates transforming growth factor-β to maintain intervertebral disc homeostasis during aging. Bone Res 6, 21 (2018).

29. Bisgrove, B. W. & Yost, H. J. The roles of cilia in developmental disorders and disease. Development 133, 4131–4143 (2006).

30. Miyamoto, T. et al. Insufficiency of ciliary cholesterol in hereditary Zellweger syndrome. EMBO J., e103499 (2020).

31. Bergmann, C., Guay-Woodford, L. M., Harris, P. C., Horie, S., Peters, D. & Torres, V. E. Polycystic kidney disease. Nat Rev Dis Primers 4, 50 (2018).

32. May-Simera, H. L. et al. Primary Cilium-Mediated Retinal Pigment Epithelium Maturation Is Disrupted in Ciliopathy Patient Cells. Cell Rep 22, 189–205 (2018).

33. Robichaux, M. A. et al. Defining the layers of a sensory cilium with STORM and cryoelectron nanoscopy. Proc. Natl. Acad. Sci. U.S.A. 116, 23562–23572 (2019).

34. Moore, E. R., Zhu, Y. X., Ryu, H. S. & Jacobs, C. R. Periosteal progenitors contribute to load-induced bone formation in adult mice and require primary cilia to sense mechanical stimulation. Stem Cell Res Ther 9, 190 (2018).

35. Yuan, X. & Yang, S. Deletion of IFT80 Impairs Epiphyseal and Articular Cartilage Formation Due to Disruption of Chondrocyte Differentiation. PLoS ONE 10, e0130618 (2015).

36. F. Fang, A. G. Schwartz, E. R. Moore, M. E. Sup, S. Thomopoulos, Primary cilia as the nexus of biophysical and hedgehog signaling at the tendon enthesis. Sci Adv 6, (2020).

37. Tang Y, Wu X, Lei W, et al. TGF-beta1-induced migration of bone mesenchymal stem cells couples bone resorption with formation. Nat Med. 2009. 15(7): 757–65.

38. Villalobos, E. et al. Fibroblast Primary Cilia Are Required for Cardiac Fibrosis. Circulation 139, 2342–2357 (2019).

39. Dalbay, M. T., Thorpe, S. D., Connelly, J. T., Chapple, J. P. & Knight, M. M. Adipogenic Differentiation of hMSCs is Mediated by Recruitment of IGF-1r Onto the Primary Cilium Associated With Cilia Elongation. Stem Cells 33, 1952–1961 (2015).

40. Pala, R., Alomari, N. & Nauli, S. M. Primary Cilium-Dependent Signaling Mechanisms. Int J Mol Sci 18, (2017).

41. Yuan, X., Serra, R. A. & Yang, S. Function and regulation of primary cilia and intraflagellar transport proteins in the skeleton. Ann. N. Y. Acad. Sci. 1335, 78–99 (2015).

42. Yuan, X. et al. Ciliary IFT80 balances canonical versus non-canonical hedgehog signalling for osteoblast differentiation. Nat Commun 7, 11024 (2016).

43. Lu, H. et al. Initiation Timing of Low-Intensity Pulsed Ultrasound Stimulation for Tendon-Bone Healing in a Rabbit Model. Am J Sports Med 44, 2706–2715 (2016).

44. Chen, C. et al. Functional decellularized fibrocartilaginous matrix graft for rotator cuff enthesis regeneration: A novel technique to avoid in-vitro loading of cells. Biomaterials 250, 119996 (2020).

45. Tang, Y. et al. Structure and ingredient-based biomimetic scaffolds combining with autologous bone marrow-derived mesenchymal stem cell sheets for bone-tendon healing. Biomaterials 241, 119837 (2020).

46. Olesen, J. L. et al. Expression of insulin-like growth factor I, insulin-like growth factor binding proteins, and collagen mRNA in mechanically loaded plantaris tendon. J. Appl. Physiol. 101, 183–188 (2006).

47. Jonsson, P., Alfredson, H., Sunding, K., Fahlström, M. & Cook, J. New regimen for eccentric calf-muscle training in patients with chronic insertional Achilles tendinopathy: results of a pilot study. Br J Sports Med 42, 746–749 (2008).

48. Ohberg, L., Lorentzon, R. & Alfredson, H. Eccentric training in patients with chronic Achilles tendinosis: normalised tendon structure and decreased thickness at follow up. Br J Sports Med 38, 8–11; discussion 11 (2004).

49. Wang, L. et al. Effects of Time to Start Training After Acute Patellar Tendon Enthesis Injuries on Healing of the Injury in a Rabbit Model. Am J Sports Med 45, 2405–2410 (2017).

50. Wada, S. et al. Postoperative Tendon Loading With Treadmill Running Delays Tendon-to-Bone Healing: Immunohistochemical Evaluation in a Murine Rotator Cuff Repair Model. J. Orthop. Res. 37, 1628–1637 (2019).

51. Zhen, G. & Cao, X. Targeting TGFβ signaling in subchondral bone and articular cartilage homeostasis. Trends Pharmacol. Sci. 35, 227–236 (2014).

52. Kim, K. K., Sheppard, D. & Chapman, H. A. TGF-β1 Signaling and Tissue Fibrosis. Cold Spring Harb Perspect Biol 10, (2018).

53. Delaney, K., Kasprzycka, P., Ciemerych, M. A. & Zimowska, M. The role of TGF-β1 during skeletal muscle regeneration. Cell Biol. Int. 41, 706–715 (2017).

54. Robertson, I. B. & Rifkin, D. B. Regulation of the Bioavailability of TGF-β and TGF-β-Related Proteins. Cold Spring Harb Perspect Biol 8, (2016).

55. Bangs, F. & Anderson, K. V. Primary Cilia and Mammalian Hedgehog Signaling. Cold Spring Harb Perspect Biol 9, (2017).

56. Hoey, D. A., Tormey, S., Ramcharan, S., O’Brien, F. J. & Jacobs, C. R. Primary cilia-mediated mechanotransduction in human mesenchymal stem cells. Stem Cells 30, 2561–2570 (2012).

57. Luo, N. et al. Primary cilia signaling mediates intraocular pressure sensation. Proc. Natl. Acad. Sci. U.S.A. 111, 12871–12876 (2014).

58. Delling, M. et al. Primary cilia are not calcium-responsive mechanosensors. Nature 531, 656–660 (2016).

59. Bell, R., Taub, P., Cagle, P., Flatow, E. L. & Andarawis-Puri, N. Development of a mouse model of supraspinatus tendon insertion site healing. J. Orthop. Res. 33, 25–32 (2015).

